# Identifying cortical substrates underlying the phenomenology of stereopsis and realness: A pilot fMRI study

**DOI:** 10.1101/557157

**Authors:** Makoto Uji, Angelika Lingnau, Ian Cavin, Dhanraj Vishwanath

## Abstract

Viewing a real scene or a stereoscopic image (e.g., 3D movies) with both eyes yields a vivid subjective impression of object solidity, tangibility, immersive negative space and sense of realness; something that is not experienced when viewing single pictures of 3D scenes normally with both eyes. This phenomenology, sometimes referred to as stereopsis, is conventionally ascribed to the derivation of depth from the differences in the two eye’s images (binocular disparity). Here we report on a pilot study designed to explore if dissociable neural activity associated with the phenomenology of realness can be localised in the cortex. In order to dissociate subjective impression from disparity processing, we capitalised on the finding that the impression of realness associated with stereoscopic viewing can also be generated when viewing a single picture of a 3D scene with one eye through an aperture. Under a blocked fMRI design, subjects viewed intact and scrambled images of natural 3-D objects and scenes under three viewing conditions: (1) single pictures viewed normally with both eyes (binocular) (2) single pictures viewed with one eye through an aperture (monocular-aperture); (3) stereoscopic anaglyph images of the same scenes viewed with both eyes (binocular stereopsis). Fixed-effects GLM contrasts aimed at isolating the phenomenology of stereopsis demonstrated a selective recruitment of similar posterior parietal regions for both monocular and binocular stereopsis conditions. Our findings provide preliminary evidence that the cortical processing underlying the subjective impression of realness may be dissociable and distinct from the derivation of depth from disparity.

## Introduction

A fundamental insight of early enquiries into visual space perception was that only certain types of visual stimulation give rise to a perceptual state in which space and objects appear visually “real” and in vivid “depth relief”. Renaissance artists correctly observed that viewing realistically rendered and optically correct perspective images (paintings) of 3D scenes produced an accurate impression of 3-dimensionality, but that they lacked the phenomenological sense of object solidity and realness that is obtained in natural scenes (Wade et al., 2001). Following Wheatstone‘s (1838) discovery of the stereoscope, contemporary understanding of this phenomenology (often referred to as stereopsis) is that it is a by-product of the derivation of depth from binocular disparities, which are the small differences between the images of the two eyes (e.g., Westheimer, 2011). In contrast to pictorial images, stereoscopic images (e.g., 3D movies) generate visual impressions of object solidity, object tangibility (the feeling that you can reach out to touch objects) and empty (negative) space between objects. Together, these impressions give the depicted objects and space a phenomenological “realness”. Motion parallax also produces a similar phenomenology which is ascribed to the fact that parallax and disparity, as sensory cues, are computationally similar (Helmholtz, 1909).

However, since the Renaissance, it has been widely observed and reported that this same phenomenology of solidity, tangibility, negative space and realness can be obtained under conditions in which neither binocular disparity nor motion parallax are present (Ames, 1925; Claparède, 1904; Koenderink, 1998; Koenderink et al., 1994; Michotte, 1948; Schlosberg, 1941; Vishwanath and Hibbard, 2013; da Vinci, cited in Wade et al., 2001; Wheatstone, 1838; Wijntjes, 2017); an observation that has been recently confirmed empirically in naïve observers, specifically when viewing single images monocularly through an aperture (Vishwanath and Hibbard, 2013). This suggests that the visual phenomenology of stereopsis is not simply tied to binocular disparity or parallax processing, but derives from a more general visual property or process. While there have been several decades of detailed investigation of the neural substrates underlying the processing of binocular disparities into depth perception (see reviews, Gonzalez and Perez, 1998; Orban, 2011; Parker, 2007; Sakata et al., 2005; Welchman, 2016), there has been no systematic investigation into the neural substrates that underlie the subjective phenomenology associated with stereopsis and visual realness. In this study, we provide an initial exploratory investigation to determine if cortical processes underlying the subjective visual phenomenology associated with real and stereoscopic 3D scenes (stereopsis) can be dissociated, localised and distinguished from disparity processing.

A large amount of work in neurophysiology has focussed on the derivation of depth and 3D structure from binocular disparity, initially in the cat and monkey, but more widely in humans in the last two decades. This work has identified substrates for initial processing of disparities, differential coding of absolute and relative disparities, differential processing of zeroth-, first- and second-order disparity relations, derivation of 3D shape from disparities, integration of disparity with other depth cues (Backus et al., 2001; Ban and Welchman, 2015; Barlow et al., 1967; Cumming, 2002; Durand et al., 2009; Georgieva et al., 2009; Goncalves et al., 2015; Neri et al., 2004; Ohzawa et al., 1990; Poggio et al., 1985; Preston et al., 2008; Taira et al., 2000; Tsao et al., 2003; Verhoef et al., 2011). These studies have shown that areas in both ventral and dorsal streams of visual processing are activated during the perception of 3D structure from disparity, but that different information is processed in each stream. While dorsal regions of the extrastriate cortex (V3A, V3B/KO, MT) and V7 have been implicated in disparity defined depth and the integration of different cues to derive the 3D structure of viewed surfaces (Backus et al., 2001; Bridge and Parker, 2007; Cottereau et al., 2011; Durand et al., 2009; Goncalves et al., 2015; Minini et al., 2010; Naganuma et al., 2005; Preston et al., 2008), ventral regions of the extrastriate cortex (i.e. V3v, V4 and LOC, PPA) have been implicated in encoding of 3D scenes, object configurations and features required for recognition and categorization (Bridge and Parker, 2007; Chandrasekaran et al., 2007; Gilaie-Dotan et al., 2002; Kourtzi and Kanwisher, 2001). Neurophysiological studies of depth processing from other cues (motion, texture and shading) are sparser but those that exist again implicate many of the same regions involved in disparity derived depth (e.g., Georgieva et al., 2008; Murray et al., 2003; Orban et al., 1999; Shikata et al., 2001; Taira et al., 2001).

Given the accumulating evidence that points to the processing of disparity and 3D structure in both ventral and dorsal visual pathways, an obvious question is whether the phenomenology of stereopsis arises in one or both visual streams, or elsewhere in the cortex. Based on the most influential theory of cortical visual processing (the two visual systems hypothesis, Goodale and Milner, 1992; Milner and Goodale, 2008), the most likely candidate for the locus of stereopsis within the visual stream is the ventral pathway, since it is implicated in conscious visual perception and recognition of objects and scenes (Bridge and Parker, 2007; Chandrasekaran et al., 2007; Gilaie-Dotan et al., 2002; Kourtzi and Kanwisher, 2001; Neri et al., 2004) while the dorsal visual pathway is generally implicated in subconscious representations guiding visuo-motor control (see reviews Anzai and DeAngelis, 2010; Freud et al., 2016; Gallivan and Culham, 2015; Goodale and Milner, 2018; Tunik et al., 2007). However, other theoretical proposals do suggest a role for the dorsal stream. For example, Michotte (1948) proposed that the phenomenology of stereopsis and realness is linked to the conscious visual awareness of the capacity to manipulate objects, implying that it arises from 3D encodings that are built up to guide manual action rather than to recognize objects. This could implicate areas of the parietal cortex involved in visuo-motor control or perhaps even other areas involved in visuo-motor planning such as the premotor (Binkofski et al., 1999; Hanakawa et al., 2008; Schubotz and Von Cramon, 2002) or prefrontal cortex (see reviews Freud et al., 2016; Gallivan and Culham, 2015; Goodale and Milner, 2018; Kravitz et al., 2011).

No prior research to our knowledge has specifically aimed to identify the substrate for the phenomenological experience of stereopsis or realness, and it is difficult to extrapolate from previous studies. The stimuli employed in most neurophysiological studies in 3D vision, where stereopsis was putatively experienced, have traditionally employed random dot (or random line) stereograms (Julesz, 1971). These types of stimuli generate an impression of 3D structure only when disparity is present and a 2D planar structure when disparity is absent (see for example: Human: Backus et al., 2001; Ban and Welchman, 2015; Goncalves et al., 2015; Neri et al., 2004; Preston et al., 2008; Monkey: Cumming, 2002; Taira et al., 2000; Tsao et al., 2003; Verhoef et al., 2010). This complicates the ability to distinguish between neural processes underlying disparity processing, representations for discrimination of depth structure or guiding movement, from those that give rise to a specific subjective experience of 3D. Interestingly, two recent studies have examined differences in cortical processing during viewing of real objects and pictures used as physical stimuli in fMRI paradigms aimed at identifying neural processes underlying object representation (Snow et al., 2011) and visually guided grasping/reaching (Freud et al., 2018). Differences in activation between real and pictured stimuli were found in both the parietal cortex (intraparietal sulcus) and the lateral occipital cortex (Freud et al., 2018; Snow et al., 2011).

In the present study, we aimed to explore the idea that the visual phenomenology of stereopsis and realness is linked to dissociable neural processes underlying 3D vision. Furthermore, we aimed to determine if this processing is distinct from activity related specifically to disparity processes. To this end, we capitalised on the fact that we could induce the phenomenology of stereopsis both in the presence of binocular disparity (binocular viewing of stereoscopic images) and in its absence (monocular-aperture viewing of single pictures). We designed stimulus and viewing conditions that were aimed at providing fMRI BOLD contrasts that would isolate potential distinctive visual processing associated with the phenomenology of realness and stereopsis under whole-brain analysis. We did not aim to establish the functional significance of any such dissociable activity (e.g., visuo-motor representations, object recognition) but simply to determine if conditions in which stereopsis is experienced has differential neural activation when other factors are controlled for.

## Methods

A 3T Siemens Trio MRI scanner with a body transmit and 32-channel receiver-array head coil was used for MR data acquisition. All data acquisition on humans was performed with approval from the relevant ethics committees (NHS Tayside and University of St. Andrews). This study was reviewed and approved by the University Teaching and Research Ethics Committee at the University of St. Andrews, and NHS Tayside. All procedures were conducted in accordance with the ethical guidelines of University of St. Andrews, NHS Tayside, and the 1964 Declaration of Helsinki. Informed consent was obtained from all participants involved in this project. Seven right-handed participants (5 males, 2 females, age = 23.8 ±7.5) took part in the study. Participants were pre-screened to confirm normal visual acuity (20/30; Snellen chart), stereonormality (Randot Test) and capacity to perceive monocular stereopsis with their right eye (abbreviated questionnaire from Vishwanath and Hibbard, 2013).

### Data acquisition

fMRI data were acquired using a GE-EPI sequence (TR=2500ms, TE=30ms, 37 slices, voxel resolution=3×3×3mm, slice gap=0.4mm, FOV = 190×190 mm, flip angle = 90°, 141 volumes). Foam padding was placed around the participant's head to reduce motion-related artefacts. A T1-weighted anatomical image (MPRAGE sequence: TR=1900ms, TE=2.64ms, 176 slices per slab, voxel resolution=0.8×0.8×1mm, GRAPPA acceleration factor=2, FOV = 200×200 mm, 9º flip angle) was also acquired.

### Stimulus display and response acquisition

Stimuli were back projected on to a screen that was built into a custom frame manufactured to fit securely into the back of the magnet bore. The screen was a Stewart Filmscreen Starglass 60 back projection screen. The screen display area was 47cm (W) x 37cm (H) at the widest dimensions. The projector was a Sanyo ET 30L with a Sanyo TNS-T11 long-throw lens running at a resolution of 1280 x 960 pixels and refresh rate of 85hz. It was located in a shielded room and the image back projected onto the screen through a wave guide. The image was viewed via a mirror (Siemens) that was attached to the head coil in front of the participants face when they lay supine on the scanner bed. The total distance to the screen from the subject’s eyes via the mirror (i.e., viewing distance) was approximately 80cm.

Three viewing conditions were tested: Monocular-aperture Pictorial (MaP), Binocular Pictorial (BP) and Stereoscopic Anaglyph (SA); see below for details. In the Binocular Pictorial condition, the eye opening of the head coil (5cm x 5cm) were both left open without any insert. For the other two conditions, custom eyepieces were constructed (plastic and cardboard) that were inserted into the head coil eye openings. For the Monocular-aperture Pictorial condition, a black eyepiece with a 7.5mm circular viewing aperture (adjustable position) was inserted into the right eye opening of the head coil, while a completely occluding black cover was inserted into the left eye opening. This left eye cover was placed over cotton and gauze that applied gentle pressure to keep the eyelid of the left eye shut. For each individual subject, before the start of the scanning, the position of the aperture in the eyepiece was adjusted so that the displayed image could be seen with the right eye but the rectangular edges of the image were occluded. For the Stereoscopic Anaglyph condition, both coil eye openings were inserted with a cover that had a 2cm x 3cm rectangular opening with thin transparent plastic filters (cyan for the right eye and red for the left eye). In both the Binocular Pictorial and Stereoscopic Anaglyph conditions, the full displayed image and screen could be seen, as well as parts of the magnet bore in the periphery. Responses (right-hand thumb press) were acquired using a Nordic Neurolabs thumb trigger response device.

### Stimuli

Stimuli (Figure 1) consisted of greyscale images of natural 3D scenes or objects that were acquired via a stereoscopic camera rig with inter-camera separation of 65mm (Hibbard, 2008). Image resolution was 800×800 pixels. There were three types of images: (1) a standard photographic (no-disparity) image, (2) the same images in a stereoscopic (red-cyan) anaglyphic format, (3) scrambled versions of the pictorial images (50×50 cells). The size of the projected images on the screen was 30cm x 30cm.

**Fig. 1.**
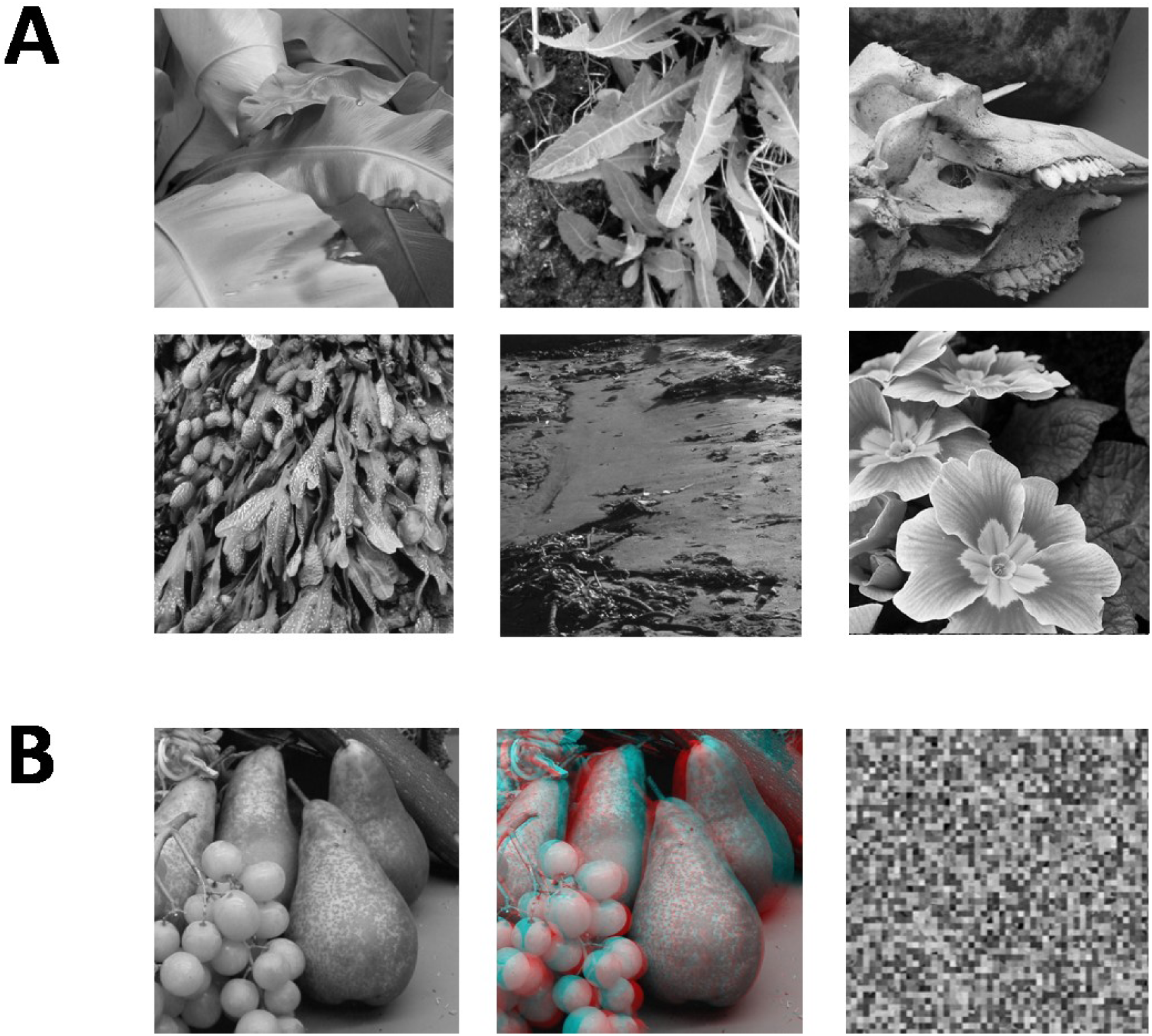
A) Sample of six of the 12 B&W photographic images used in the study. B) Example of the three versions of each image that were used in the experiment. The leftmost image is the pictorial (no-disparity) version of the image (left-camera view only). The middle image is the stereoscopic (disparity) version of the image combining left and right camera views. The rightmost image is the scrambled version of the pictorial image (50×50 cells)

### Viewing conditions

Participants viewed visual stimuli under three types of viewing conditions, which were each tested on separate runs, with bed retractions as necessary between viewing conditions to change the eyepieces. The three viewing conditions were Binocular Pictorial: viewing a pictorial (no disparity) image or a scrambled version naturally through both eyes without visual restriction. Monocular-aperture Pictorial: viewing a pictorial (no disparity) image or a scrambled version with the right eye only through a circular aperture that occluded the rectangular edges of the images; Stereoscopic Anaglyph: viewing either a stereoscopic image (disparity) or pictorial version of the same image (no disparity) through red-cyan anaglyph filters.

### fMRI design

We used a blocked fMRI design (see Figure 2). Each block began with a pre-stimulus fixation phase of 7.5 seconds, followed by the sequential display of 10 images in random order (2000ms per image) for a total of 20 seconds. For the MaP and BP conditions, there were 12 blocks per run alternating between intact pictorial images (6 blocks) and scrambled images (6 blocks). For the SA condition, there were also 12 blocks per run, alternating between stereoscopic images (disparity, 6 Blocks) and pictorial images (no disparity, 6 blocks). The start of each run was preceded by a 10 second fixation phase. For each viewing condition, we acquired data for two consecutive 12-block runs, and the bed was retracted partially in order to change the eyepieces for a subsequent viewing condition. The order of acquisition for the three types of runs (MaP, BP, SA) were partially counterbalanced across participants. The anatomical image was acquired in between two of different viewing condition runs so as to provide a period of task-free rest.

**Fig. 2.**
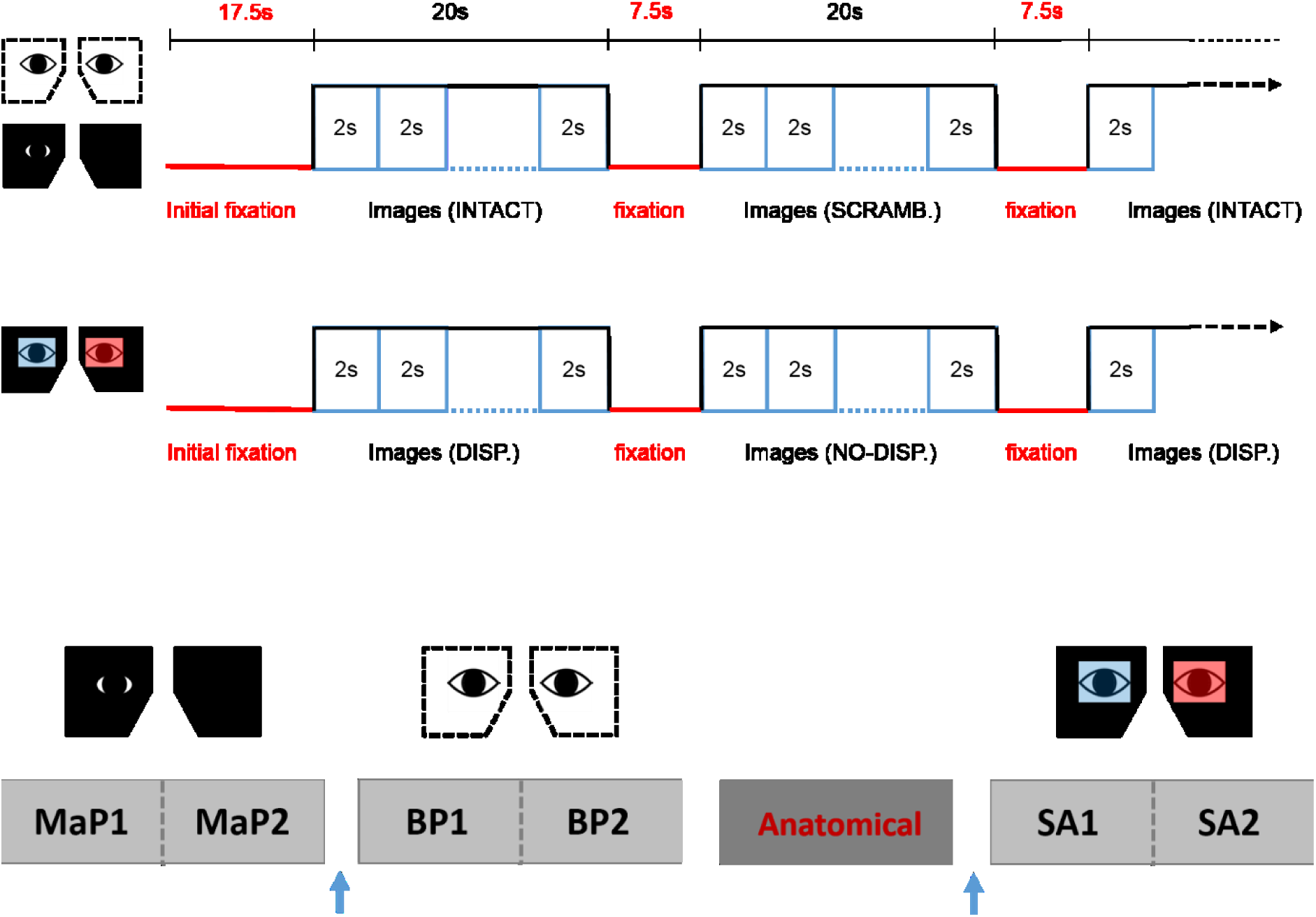
fMRI design. A) The upper panel shows the stimulus sequence for the Binocular Pictorial (BP) and Monocular-aperture Pictorial (MaP) conditions, while the lower panel shows the sequence for the Stereoscopic Anaglyph (SA) condition. B) An example sequence of the functional runs and anatomical scans for one participant. Two runs for each viewing condition were done consecutively with a brief break in between. The order of the viewing conditions and the anatomical scan was partially counterbalanced across participants. The blue arrow represents partial bed retractions to change eyepieces.

### Visual fixation task

In order to control fixation and attention across all viewing conditions and stimulus types, participants were required to do the same visual detection task at central fixation for all runs while passively viewing the displayed images. Participants were instructed to keep their eyes on the fixation dot (a 4mm x 4mm square), monitor its luminance and press the response button as quickly as possible (right hand thumb trigger) whenever it turned from black to grey. The luminance change occurred at random intervals during the stimulus blocks. Specifically, the luminance change occurred in 50% of randomly selected individual stimulus trials within a block, 1000ms after stimulus onset. Participants were instructed to remain as still as possible during the entire duration of the experiment and also during bed-retractions.

#### Analysis

fMRI data were processed using FSL v5.0.9 (https://fsl.fmrib.ox.ac.uk/fsl/). Data from each participant were motion-corrected (MCFLIRT), spatially smoothed (5mm FWHM Gaussian kernel), high-pass temporally filtered (100s cutoff), registered to their T1 anatomical brain image (FLIRT), and normalised to the MNI 2mm standard brain. GLM analyses were performed using FEAT v6.0. First-level analysis was performed employing two regressors (time-locked to the onset of each 20 sec block): 1) intact stimuli, 2) scrambled stimuli for the MaP and BP; and 1) disparity stimuli, 2) non-disparity stimuli for the SA. All regressors were convolved with a double-gamma HRF.

For the Monocular-aperture Pictorial (MaP) and Binocular Pictorial (BP) conditions, we computed the first-level contrasts (intact > scrambled stimuli), whereas for the Stereoscopic Anaglyph (SA) condition, we computed the first-level contrast (disparity > non-disparity stimuli). For each participant, the first-level results were combined across the two runs of each viewing condition using a second-level, fixed effects analysis to calculate an average response per subject for each viewing condition. To determine areas recruited during the processing of monocular stereopsis, we computed the contrast [MaP_INTACT_> MaP_SCRAMB_] > [BP_INTACT_> BP_SCRAMB_] at the third-level, fixed effects analysis per participant. These second and third level results were then combined across all participants at the fourth, group-level using a FLAME fixed-effects analysis (Woolrich et al., 2004) for each viewing condition and monocular stereopsis contrast. The critical contrasts were the fourth-level fixed effects results for the monocular stereopsis [MaP_INTACT_> MaP_SCRAMB_] > [BP_INTACT_> BP_SCRAMB_], and the fourth-level fixed effects results for the binocular stereopsis [SA_DISP_ > SA_NO-DISP_]. Since we did not expect to find any results related to the processing of stereopsis outside grey matter based on our a-priori hypothesis, we applied a mask of grey matter (FSL FAST, https://fsl.fmrib.ox.ac.uk/fsl/) (Zhang et al., 2001) as pre-threshold mask to all group-level statistical maps (Eickhoff et al., 2009; Uji et al., 2018). Main effect (boxcar model of the task) BOLD Z-statistic images were thresholded at uncorrected significance level of p < 0.001 (Eklund et al., 2016; Woo et al., 2014).

We also conducted a conjunction analysis (Friston et al., 2005; Nichols et al., 2005) to obtain the commonly activating brain regions between the effects of monocular and binocular stereopsis. The two Z-statistic images from each effect were tested against the conjunction null hypothesis at uncorrected significance level of p < 0.001.

Due to the limitations of voxelwise analysis and increased risk of Type I error using uncorrected (p<0.001) thresholds for multiple comparisons, we applied the false discovery rate (FDR) measure (Benjamini and Hochberg, 1995) using the FDR analysis tool supplied by the FSL software (Jenkinson et al., 2012; Nichols, 2012). FDR represents the expected proportion of rejected hypotheses that are false positives. To correct for the multiple comparisons, the two p-maps from the binocular and monocular stereopsis contrasts were further thresholded to a FDR of 5%.

## Results

All participants performed the visual detection task at fixation as instructed showing a group mean accuracy of 98.8 ± 0.3 % (± standard error [SE]) for the MaP condition, 98.8 ± 0.4 % for the BP condition, and 94.6 ± 3.8 % of the trials for the SA condition. Accuracy did not differ significantly between viewing conditions, *F*(2,12)=1.34, *p*=0.30.

We first examined areas revealed by the contrasts [MaP_INTACT_> MaP_SCRAMB_] and [BP_INTACT_> BP_SCRAMB_]. These contrasts aimed to identify activations associated with perception of objects, scenes and 3D structure under each of the two viewing conditions (e.g., Epstein and Kanwisher, 1998; Kourtzi and Kanwisher, 2001). Both contrasts revealed similar statistical maps (z-score > 3.1, p<0.001, fixed effects) that included regions corresponding to both dorsal and ventral aspects of higher occipital areas, parietal cortex and posterior aspects of the cingulate cortex (Figure 3 and 4). Peak responses for the monocular aperture condition were found in the left parietal cortex [peak voxel: Z=11.2, p<1×10^−16^, MNI coordinates (x, y, z): −32, −72, 38], in the right parietal cortex [peak voxel: Z=7.99, p<1×10^−15^, MNI coordinates: 20, −82, 42], and in the left and right posterior cingulate [peak voxels: Z=6.63, p<1×10^−10^ and Z=6.44, p<1×10^−10^, MNI coordinates: −26, −56, 10 and 28, −58, 4] respectively. Peak responses for the binocular pictorial condition were found in the left parietal cortex [peak voxel: Z=6.34, p<1×10^−9^, MNI coordinates: −6, −80, 44], in the right parietal cortex [peak voxels: Z=7.05, p<1×10^−12^ and Z=5.47, p<1×10^−7^, MNI coordinates: 44, −76, 20 and 46, −54, 46] respectively, in the left posterior cingulate [peak voxels: Z=6.76, p<1×10^−11^ and Z=4.49, p<1×10^−5^, MNI coordinates: −26, −54, 4 and −4, −34, 24] respectively, in the right posterior cingulate [peak voxel: Z=6.61, p<1×10^−10^, MNI coordinates: 28, −52, 6], and in the left lateral occipital cortex [peak voxel: Z=6.43, p<1×10^−10^, MNI coordinates: −38, −60, −6].

**Fig. 3.**
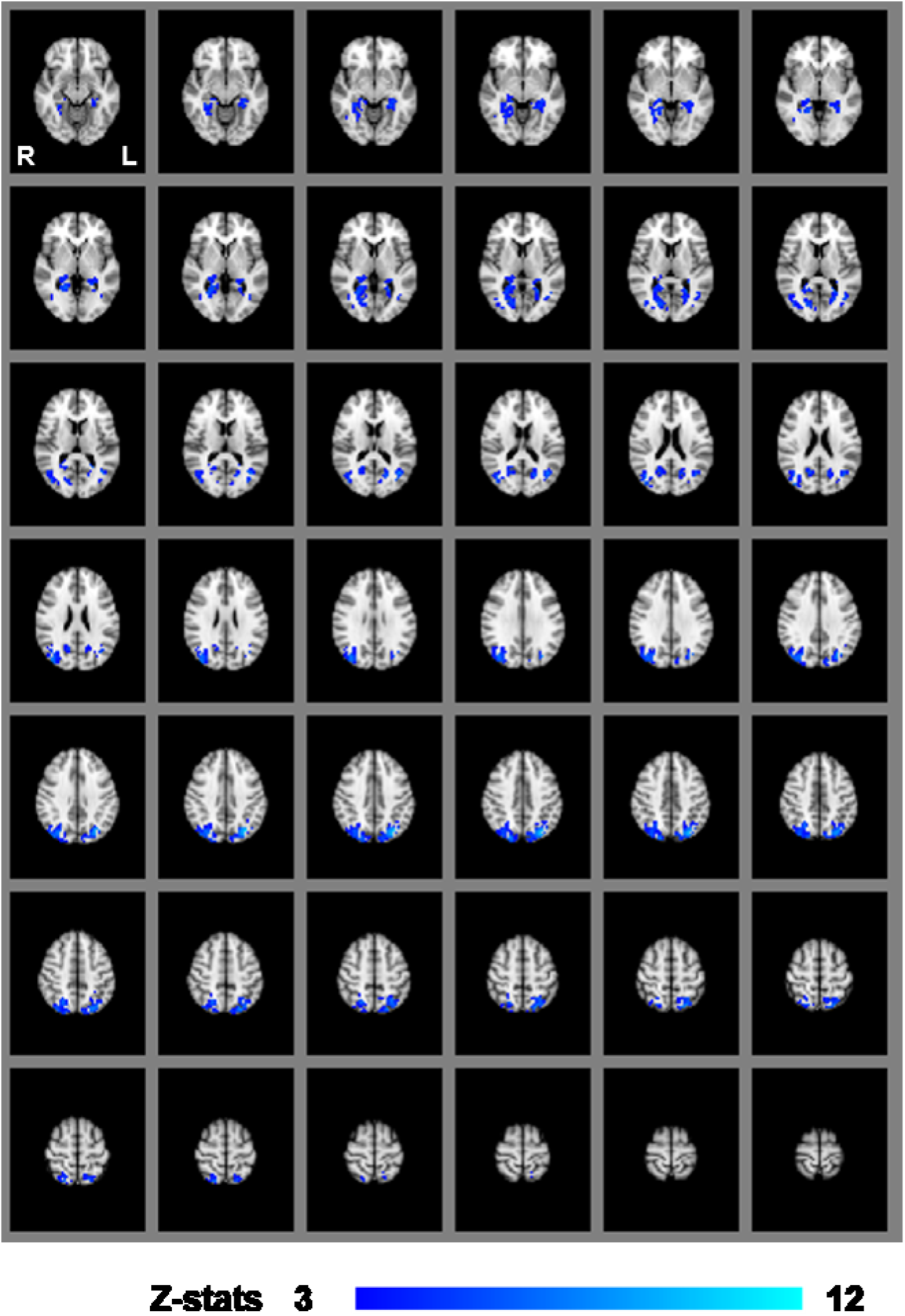
Group results for monocular-aperture viewing, revealed by the fixed-effects (N = 7) GLM contrast [MaP_INTACT_ > MaP_SCRAMB_ images]. All responses are masked to grey matter. Uncorrected p < 0.001.

**Fig. 4.**
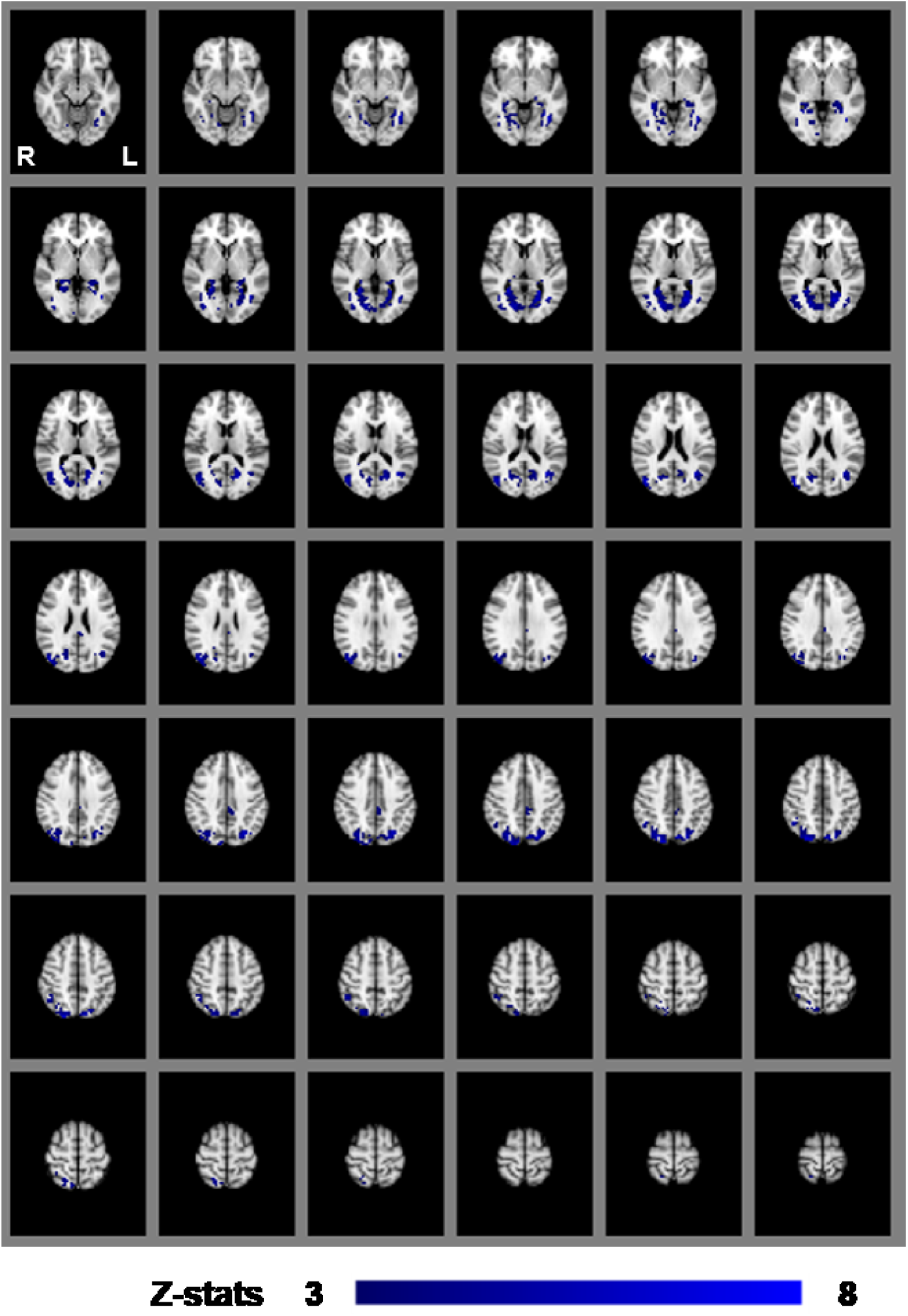
Group results for binocular viewing, revealed by the fixed-effects (N=7) GLM contrast [BP_INTACT_ > BP_SCRAMB_ images]. All responses are masked to grey matter. Uncorrected p < 0.001.

To identify areas contributing to the impression of monocular stereopsis, we examined the contrast ([MaP_INTACT_> MaP_SCRAMB_] > [BP_INTACT_> BP_SCRAMB_]). This contrast revealed a cluster in the left parietal cortex [peak voxel: Z=5.30, p<1×10^−7^, MNI coordinates: −28, −60, 54] and in the right parietal cortex [peak voxels: Z=5.28, p<1×10^−7^ and Z=4.01, p<1×10^−4^, MNI coordinates: 36, −74, 50 and 20, −64, 50] (Figure 5).

**Fig. 5.**
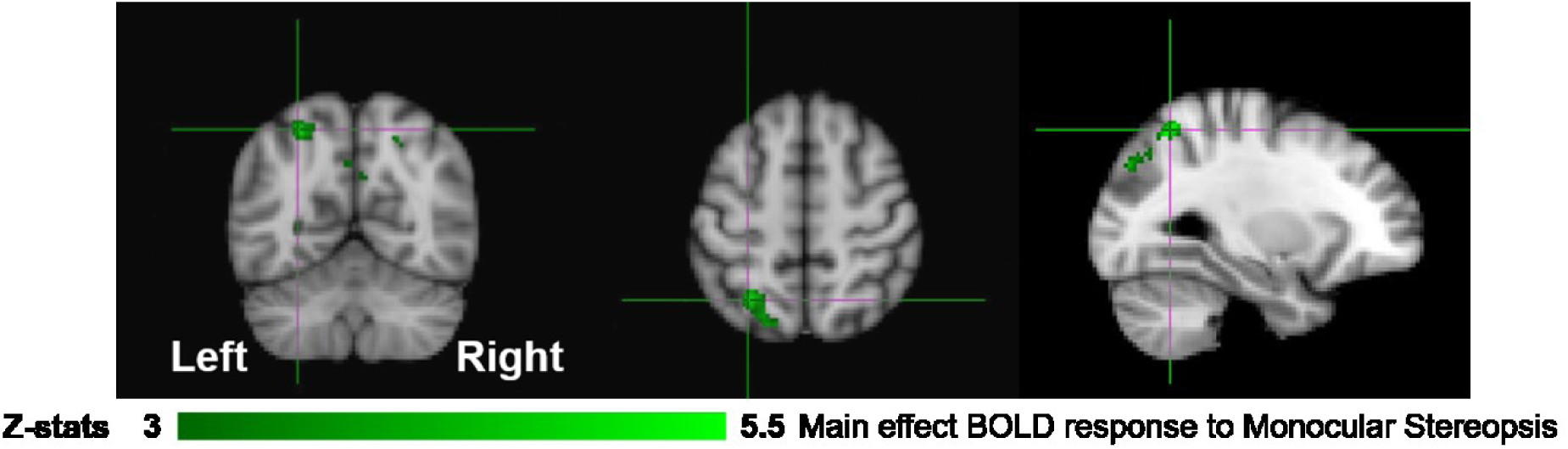
Group results for monocular stereopsis, revealed by the fixed-effects (N=7) GLM contrast [MaP_INTACT_> MaP_SCRAMB_] > [BP_INTACT_> BP_SCRAMB_]. All responses are masked to grey matter. Uncorrected p < 0.001. The crosshairs represent the peak in the left parietal cortex [MNI coordinates: −28, −60, 54].

To identify areas that are involved both in the processing of disparity and binocular stereopsis, we used the contrast [SA_DISP_ > SA_NO-DISP_]. This contrast revealed a cluster in the left parietal cortex [peak voxels: Z=3.87, p<1×10^−4^ and Z=4.18, p<1×10^−4^, MNI coordinates: −28, −74, 46 and −30, −44, 50], in the left lateral occipital cortex [peak voxel: Z=3.66, p<0.001, MNI coordinates: −26, −84, 18], and in the right lateral occipital cortex [peak voxels: Z=4.11, p<1×10^−4^ and Z=4.16, p<1×10^−4^, MNI coordinates: 36, −76, 4 and 42, −76, 26] (Figure 6).

**Fig. 6.**
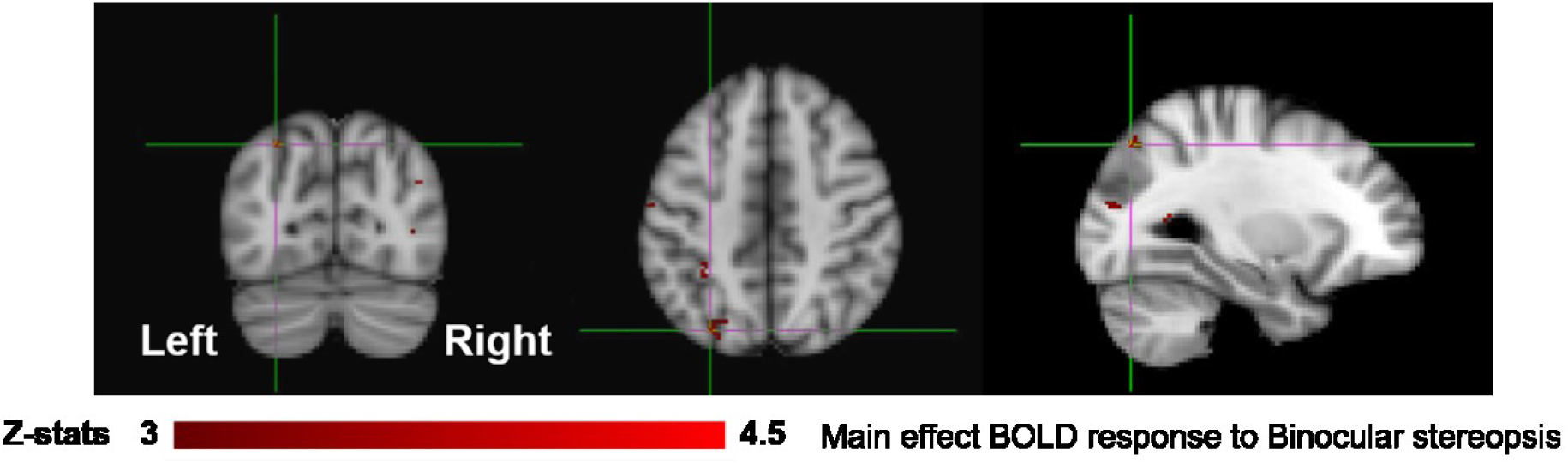
Group results for binocular stereopsis, revealed by the fixed-effects (N=7) GLM contrast [SA_DISP_ > SA_NO-DISP_]. All responses are masked to grey matter. Uncorrected p < 0.001. The crosshairs represent the peak in the left parietal cortex [MNI coordinates: −28, −74, 46].

Figure 7 shows the overlay of regions of significant activation for both the monocular stereopsis and binocular stereopsis contrasts. In order to identify potential activity associated with the phenomenology of stereopsis distinct from activity arising from disparity processing we conducted a conjunction analysis to determine regions that were significantly overlapping between the monocular and binocular stereopsis contrasts. The analysis confirmed a region in the left parietal cortex at [−24, −76, 46] mm *[MNI:x,y,z]* as significantly (p<0.001).

**Fig. 7.**
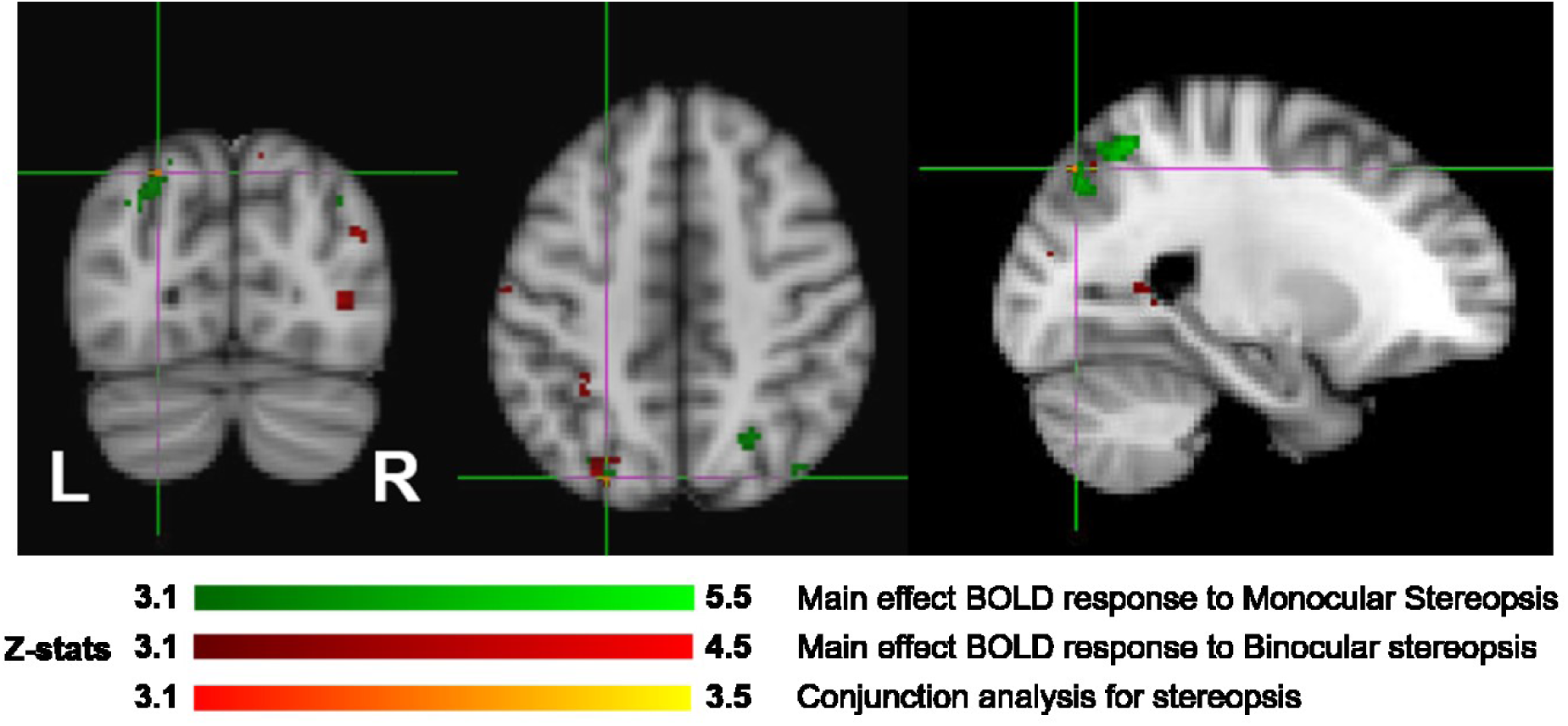
Group results (N=7) for monocular (green) and binocular (red) stereopsis, and conjunction analysis for monocular and binocular stereopsis (red-yellow), revealed by the fMRI fixed effects results. The main effects are masked to grey matter. Uncorrected p < 0.001. The crosshairs represent the significantly overlapping region between monocular and binocular stereopsis [MNI coordinates: −24, −76, 46].

Due to the probability of type-1 errors in the uncorrected voxelwise analysis, we applied a false discovery rate (FDR) of 0.05 to the uncorrected (P<0.001) thresholds. The FDR corrected p <0.001 threshold revealed significant clusters only in the monocular stereopsis contrast and not in the binocular stereopsis contrast. Specifically, in the monocular stereopsis contrast two clusters in the left parietal cortex [MNI coordinates: −28, −60, 54 and −28, −80, 36] and a cluster in the right parietal cortex [MNI coordinates: 36, −74, 50] were identified (Figure 8).

**Fig. 8.**
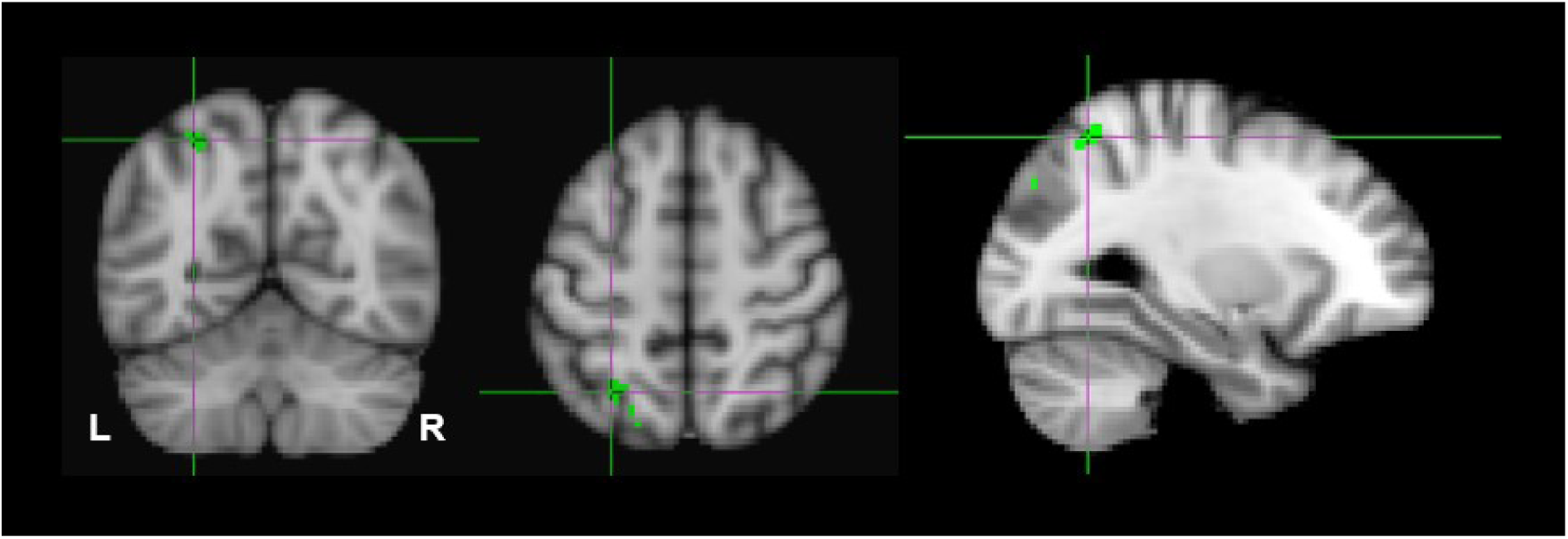
Group results for monocular stereopsis, revealed by the fixed-effects (N=7) GLM contrast [MaP_INTACT_> MaP_SCRAMB_] > [BP_INTACT_> BP_SCRAMB_]. All responses are masked to grey matter. FDR (p < 0.05) corrected p < 0.001. The crosshairs represent the peak in the left parietal cortex [MNI coordinates: −28, −60, 54].

## Discussion

The sensation of realness associated with binocular stereopsis is one of the central attributes of human perceptual experience of the visual world and has been the subject of enquiry since the very beginning of empirical science. Our exploratory study developed and tested an experimental paradigm aimed at identifying the potential neural underpinnings of this important visual attribute. Our paradigm capitalised on findings that this sensation can be produced under significantly different viewing and stimulus condition (stereoscopic images and monocular-aperture viewing of single pictorial images). This allowed us to attempt to distinguish experimentally between the processing of sensory information (binocular disparity) that is conventionally believed to give rise to the phenomenology and the visual phenomenology itself.

Specifically, we measured the amplitude of the BOLD signal while participants passively viewed images of natural scenes or objects under three different viewing conditions (Binocular Pictorial: BP, Stereoscopic Anaglyph: SA, Monocular-aperture Pictorial: MaP). Both the MaP and SA (disparity) conditions were expected to induce the qualitative impressions of stereopsis and realness. We hypothesised that conditions yielding binocular and monocular stereopsis would reveal both different and similar clusters of voxels: different because the source of signals specifying 3D structure differ in the two cases; similar because in both cases the same 3D objects and scenes are viewed and perceived. We also reasoned that the existence of a generic visual process or attribute related to the phenomenology of stereopsis (and any associated cognitive state) should imply unique dissociable loci of neural processing regardless of the source of sensory signals. On this interpretation, we expected that we would find specific overlapping cortical regions recruited in contrasts aimed at titrating signals linked to binocular and monocular stereopsis. Since we had no a-priori region of interest in the cortex, expecting potential substrates in visual, parietal, temporal, motor, premotor, or prefrontal cortices we conducted a whole-brain analysis.

In order to dissociate activity specifically linked to the perception of objects, scenes and 3D structure, we examined fixed-effects GLM contrasts under both binocular and monocular-aperture viewing of single pictures (BP_INTACT_>BP_SCRAMB_ and MaP_INTACT_>MaP_SCRAMB_). As expected, we found generally similar results in these two contrasts, with significant clusters in both dorsal and ventral aspects of the visual stream as well as cingulate cortex (Figures 3 and 4), consistent with areas typically identified with the perception of objects, scenes and 3D structure (Denys et al., 2004; Epstein and Kanwisher, 1998; Georgieva et al., 2008; Kourtzi and Kanwisher, 2001; also see a review Orban et al., 2004). In order to specifically dissociate activity linked to the phenomenology of realness and stereopsis from the generic perception of objects, scenes and 3D structure, we examined the contrasts of monocular stereopsis ([MaP_INTACT_>MaP_SCRAMB_] > [BP_INTACT_>BP_SCRAMB_]) and binocular stereopsis (SA_DISP_>SA_NO-DISP_). For the binocular stereopsis contrasts (SA_DISP_>SA_NO-DISP_; Figure 6) we identified significant clusters in left posterior parietal cortex (PPC), and bilateral lateral occipital cortex (LOC), in agreement with previous human and non-human primate fMRI studies implicating the processing of binocular disparity into 3D perception in both ventral and dorsal aspects of the visual stream (LOC: Bridge and Parker, 2007; Chandrasekaran et al., 2007; Gilaie-Dotan et al., 2002; Kourtzi and Kanwisher, 2001; Neri et al., 2004; PPC: Durand et al., 2009, 2007; Georgieva et al., 2009; Joly et al., 2009; Minini et al., 2010; Rosenberg et al., 2013; Rosenberg and Angelaki, 2014; Taira et al., 2000; Tsao et al., 2003; Tsutsui et al., 2002; Van Dromme et al., 2016, 2015; Verhoef et al., 2015). For the monocular stereopsis contrast ([MaP_INTACT_>MaP_SCRAMB_] > [BP_INTACT_>BP_SCRAMB_]; Figure 5), we identified significant clusters in bilateral PPC without the recruitment of LOC. While the differences between the two stereopsis contrasts may be attributable to differences in the construction of the contrasts themselves (namely, derived from a third vs. a second level contrast, respectively), the processed signals (disparity is present in the stimulus in the binocular stereopsis contrast but not in the monocular stereopsis contrast) did also drive the differences. Finally, a conjunction analysis between the monocular and binocular stereopsis contrasts (Figure 7) identified significant voxels only in left posterior parietal cortex at [−24, −76, 46] mm [MNI:x,y,z] which is putative posterior intraparietal sulcus (IPS) (Durand et al., 2009; Georgieva et al., 2009; Zlatkina and Petrides, 2014). Previous research has implicated the IPS in binocular disparity defined depth (Non-human primate fMRI: Durand et al., 2007; Joly et al., 2009; Rosenberg et al., 2013; Rosenberg and Angelaki, 2014; Taira et al., 2000; Tsao et al., 2003; Tsutsui et al., 2002; Van Dromme et al., 2016, 2015; Verhoef et al., 2015; Human fMRI: Chandrasekaran et al., 2007; Durand et al., 2009; Georgieva et al., 2009; Minini et al., 2010; Tsao et al., 2003) and also the visuo-motor transformations and visual localization in 3D space that permits guided motor control and interaction (Buchwald et al., 2018; Chen et al., 2018; Freud et al., 2016; Gallivan and Culham, 2015; Grefkes and Fink, 2005; Konen et al., 2013; Tunik et al., 2007; Vingerhoets, 2014).

These results provide the first glimpse that the neural processing specifically associated with the phenomenology of visual realness (stereopsis) can be dissociated. Moreover, the results revealed that binocular and monocular stereopsis share underlying neural substrates. From a theoretical standpoint, the findings of dissociable activity in posterior parietal cortex regions implicated in visuo-motor control (IPS) lend support to the claim that the visual impression of realness is associated with the conscious awareness of the capacity to manipulate 3D objects (Michotte, 1948). The results are also consistent with other recent work in fMRI that has implicated similar regions of the parietal cortex in differentiating visual perception guiding movements to either real or pictured objects (Freud et al., 2018) and our own recent EEG data revealing dissociable EEG gamma activity in the posterior parietal cortex associated with the qualitative impression of monocular stereopsis (Uji et al., under review).

### Limitations

The exploratory nature of this study presents several limitations to the interpretation of our results. First and foremost is that our sample size was limited to seven participants due to the restricted scanner time that was funded for this project^1^. Ideally, for a whole-brain analysis with no a-priori ROIs, we would have preferred to test 15-20 participants. Due to the limited sample size, we reasoned that we would not have the power to identify dissociative activations based on standard random-effects GLM analyses. Therefore, we used a FLAME fixed-effect analysis at the fourth group-level analysis averaging across all participants (Matthews and Jezzard, 2004; Woolrich et al., 2004). The fixed effect analysis provides an expression of changes in the group mean signal relative to the group pooled within-subject variance, not taking into any consideration of any intersubjective variability. This makes our findings hard to generalize to a larger population outside the sampled group (Friston et al., 2005; Matthews and Jezzard, 2004). Furthermore, in fixed effect analyses there is high risk that the results are driven primarily by one or two participants only. This could indeed be an interpretation of our results as individual subject z-scores of the monocular stereopsis contrast show that significant level of the z-scores were only seen for two or three participants at the group-wise peak voxel locations identified by the conjunction analysis (see Figure 9). More precisely, for the monocular stereopsis contrast, which was the only contrast where there were surviving clusters after FDR corrections at the group level, it appears that subjects 1 and 4 predominantly contributed to the group level significant clusters in the parietal cortex. Nonetheless, five out of seven participants revealed similar BOLD responses in those significant regions, thus resulting in the fixed-effect FDR corrected significant clusters at the group level (see Figure 9).

**Fig. 9.**
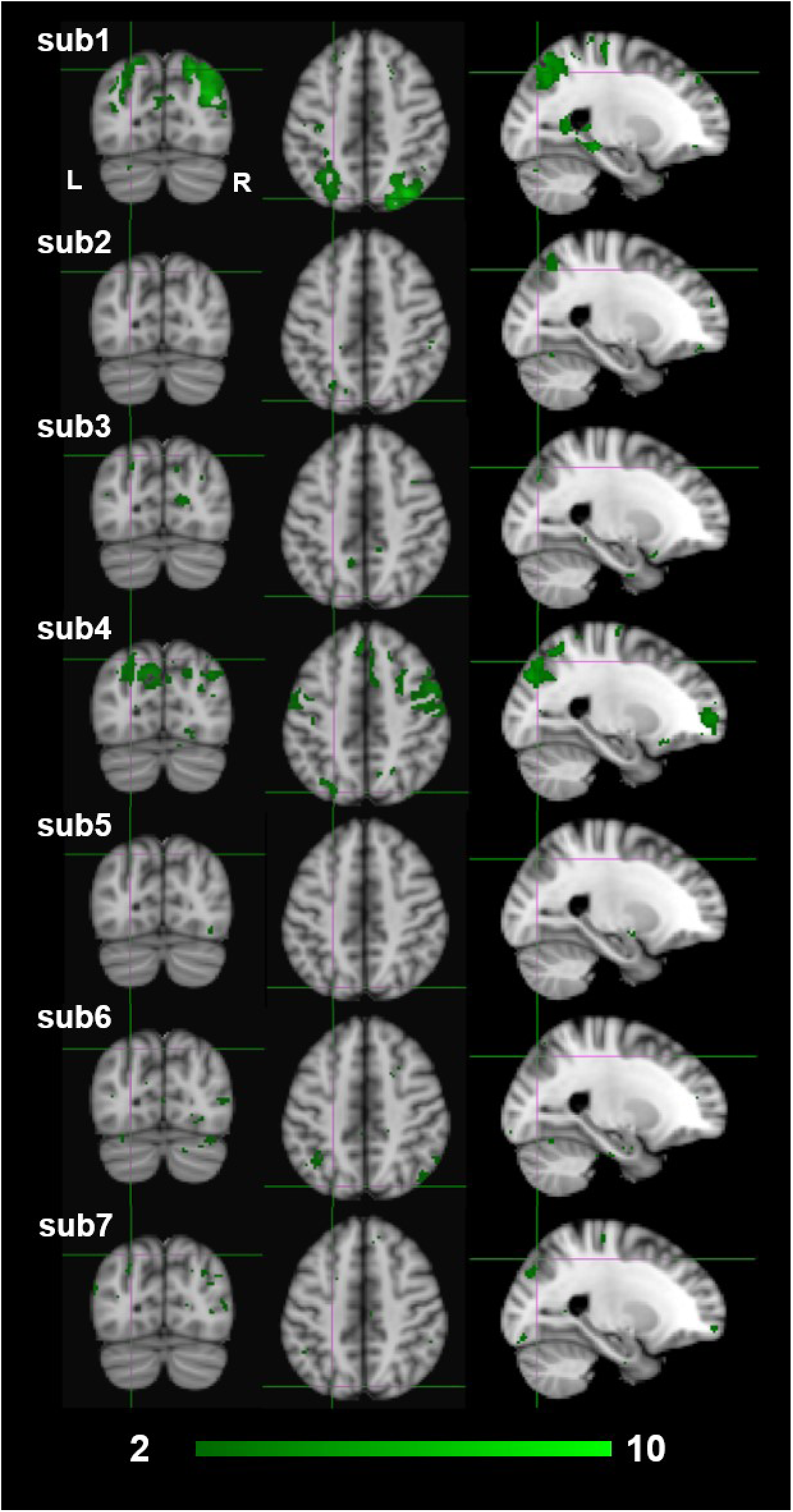
Individual participants results by the third-level fixed effects contrast ([MaP_INTACT_ > MaP_SCRAMB_] > [BP_INTACT_> BP_SCRAMB_]) for monocular (green) stereopsis. The crosshairs represent the significantly overlapping region between monocular and binocular stereopsis [MNI coordinates: −24 −76 46].

Second, due to the power limitations, we initially did not employ any family wise error correction (e.g., Bonferroni, False Discovery Rate), or apply the more conventional cluster-based thresholding approach (Eklund et al., 2018, 2016; Woo et al., 2014), thresholding-free cluster enhancement (TFCE) approach (Smith and Nichols, 2009), or randomisation methods/non-parametric permutation methods (Winkler et al., 2014). Instead, we opted for a simple voxel-based thresholding at uncorrected p<0.001 instead of more liberal approach of a cluster-based thresholding using Z > 2.3 for multiple comparisons (Bansal and Peterson, 2018; Eklund et al., 2018, 2016; Woo et al., 2014). We are aware of the risks this raises of false positives in our findings. Therefore, we applied False Discovery Rate (FDR) corrections (p<0.05) on both binocular and monocular stereopsis contrasts (Benjamini and Hochberg, 1995; Nichols, 2012). Applying FDR corrections revealed no significant clusters in the binocular stereopsis contrast, whereas the monocular stereopsis contrast revealed two significant clusters in the left parietal cortex and one significant cluster in the right parietal cortex (see Figure 8).

Third, our goals here were quite modest, to simply see if under near whole-brain analysis (excluding the cerebellum), we could identify voxels that dissociate between conditions in which stereopsis and the phenomenology or realness is present or absent. We did not aim to test specific hypotheses based on a-priori ROIs or establish what such dissociable activity might entail (e.g., visuo-motor representations, recognition, cognitive states). Previous fMRI studies on 3D vision have typically limited analysis to the visual cortex and regions of extra striate visual pathways as well as measuring retinotopic mapping in order to increase spatial resolution and localization in the visual cortex (Backus et al., 2001; Ban and Welchman, 2015; Goncalves et al., 2015; Neri et al., 2004). Similarly, studies in visuo-motor control have also limited coverage over the visual, parietal, and motor cortex (Culham et al., 2003; Gallivan et al., 2011, 2009)^2^. Future investigation of the phenomenology of stereopsis and realness would likely yield more robust results taking this more fine-grained approach, along with increased sample size both at the participants and runs/trials level and employment of multivoxel pattern analysis methods to analyse distributed patterns of neural activity to infer the functional role of brain areas and networks engaged during this phenomenology (Haynes, 2015; Norman et al., 2006).

### Implications: Functional links to depth processing

Interestingly, recent studies have examined differences in the cortical processing of visual stimulation arising from real or pictured objects. The focus of these studies was on the difference between real objects and pictures as physical stimuli in fMRI paradigms aimed at identifying neural processes underlying object representation (Snow et al., 2011) and visually guided grasping/reaching (Freud et al., 2018). Specifically, Snow et al. (2011) found that regions of the intraparietal sulcus and lateral occipital cortex, which showed typical repetition adaptation effects to pictured objects, did not show such effects for real objects. Freud et al. (2018) similarly found differential activation in the intraparietal sulcus to real 3D objects vs. 2D images during the planning phase (pre-movement initiation) of grasping and reaching motor tasks. Although these studies did not aim to specifically identify which visual attributes, cues or visual processing underlie the difference between real and pictured objects, their findings do provide support for our overall interpretation that dissociable patterns of results exists that distinguishes between the phenomenology of real depth (stereopsis) and pictorial depth.

How can this be linked to existing and future work in the neurophysiology of depth perception? In most observers, binocular disparity produces the strongest impression of the phenomenological attributes of stereopsis (object solidity, negative space, tangibility). Several intraparietal sulcus regions (dorsal IPS anterior, dorsal IPS medial, parieto-occipital IPS and ventral IPS regions) have been shown to be recruited in the presence of binocular disparity defined depth structure (Durand et al., 2009; Georgieva et al., 2009). For example, Durand et al. (2009) demonstrated that human anterior intraparietal sulcus (Dorsal IPS medial at [−22, −62, 56] mm [MNI:x,y,z]; dorsal IPS anterior at [−30, −50, 64] mm) was recruited when processing 3D structure, whereas the posterior IPS [−24, −82, 32] mm was processed when processing location in 3D space. This shows that disparity processing occurs along the dorsal visual pathway all the way through the regions encoding responsible for visually guided manual action encoding both location in depth and 3D shape. Visual guidance of movement, in principle, requires the derivation of absolute values of object dimensions scaled to the motor plant (e.g. separation of thumb and fingers for grasping; required distance of hand transport for reaching; Kopiske et al., 2018; Glennerster et al., 1996); on the other hand, the 3D shape of an object can be specified fully by only relative depth values. How might this be linked to the difference between the phenomenology of stereopsis and perception of pictorial depth? A recent view is that the primary phenomenological components of the experience of stereopsis (the impression of tangibility, solidity and negative space) is linked to successful encoding of scaled intra- and inter-objects dimensions required for visuo-motor control, while pictorial depth (no stereopsis) constitutes 3D perception without scale (Vishwanath, 2014). Since binocular disparity is widely considered the best signal for the derivation of scaled depth values (Watt and Bradshaw, 2003), the conjecture can explain why the phenomenology of stereopsis and realness is most vividly seen in the presence of binocular disparity, but like Michotte’s proposal, it does not restrict the phenomenology of stereopsis to conditions where only binocular disparity or motion parallax is present. No studies to our knowledge have examined information processing along the dorsal stream in terms of depth or disparity scaling, likely because it is difficult to design parametric stimulus sets that dissociate changes in this attribute (depth scaling) from changes in other variables, such as retinal size, stimulus size, stimulus distance, disparity magnitude and perceived depth magnitude. Genuinely integrating questions arising from visual phenomenology and visual function (depth discrimination, estimation, visuo-motor capacity) via theoretically motivated models may begin to yield new ways in which to interrogate and operationalize these questions

Here we have reported on the first attempt at just such an integration utilizing a neuroimaging paradigm. Future neuroimaging research will need to provide more robust confirmatory evidence and determine if the representation and processes that bring about the experience of stereopsis and realness are the same regardless of the source of the depth signal (binocular disparity, motion parallax, synopter or pictorial cues) and also identify the specific stage of transformation of visual information that underlies this central phenomenological aspect of human 3D space perception.

## Acknowledgements

Support for DV and MU was provided by the Leverhulme Trust.

1 This project was conducted under a research development budget awarded by Clinical Research Centre, NHS Tayside.

